# Embodiment of sleep-related words: evidence from event-related potentials

**DOI:** 10.1101/2020.12.23.424194

**Authors:** Mareike J. Hülsemann, Björn Rasch

**Affiliations:** University of Fribourg, Department of Psychology, Division of Cognitive Biopsychology and Methods, Rue P.A. de Faucigny 2, 1701 Fribourg, Switzerland

**Keywords:** N400, embodied cognition, multimodal representation, language perception, auditory word categorisation

## Abstract

Our thoughts, plans and intentions can influence physiological sleep, but the underlying mechanisms are unknown. According to the theoretical framework of “embodied cognition”, the semantic content of cognitive processes is represented by multimodal networks in the brain which also include body-related functions. Such multimodal representation could offer a mechanism which explains mutual influences between cognition and sleep. In the current study we tested whether sleep-related words are represented in multimodal networks by examining the effect of congruent vs. incongruent body positions on word processing during wakefulness.

We experimentally manipulated the body position of 66 subjects (50 females, 16 males, 19-40 years old) between standing upright and lying down. Sleep- and activity-related words were presented around the individual speech recognition threshold to increase task difficulty. Our results show that word processing is facilitated in congruent body positions (sleep words: lying down and activity words: standing upright) compared with incongruent body positions, as indicated by a reduced N400 of the event-related potential (ERP) in the congruent condition with the lowest volume. In addition, early sensory components of the ERP (N180 and P280) were enhanced, suggesting that words were also acoustically better understood when the body position was congruent with the semantic meaning of the word. However, the difference in ERPs did not translate to differences on a behavioural level.

Our results support the prediction of embodied processing of sleep- and activity-related words. Body position potentially induces a pre-activation of multimodal networks, thereby enhancing the access to the semantic concepts of words related to current the body position. The mutual link between semantic meaning and body-related function could be a key element in explaining influences of cognitive processing on sleep.

## 1. Introduction

Our thoughts affect our sleep. Cognitive processes such as rumination and worries are key aspects underlying insomnia development and maintenance (Harvey et al., 2005; Riemann et al., 2010). Effective psychotherapeutic interventions to treat insomnia typically try to reduce, change or replace these types of “cognitive arousal”. In addition, “calming” of thoughts using diverse relaxation techniques can have sleep promoting effects (Riemann et al., 2017). In particular, through a series of studies we have shown that verbal suggestions to sleep deeper given before sleep can extend the time spent in objectively measured slow-wave sleep (SWS) (Cordi et al., 2014; Cordi et al., 2015; Cordi et al., 2020). However, the underlying mechanisms by which our thoughts or verbal suggestions influence neurophysiological sleep regulation are still unknown.

We have previously proposed that certain cognitive processes could remain active during sleep when initiated before sleep (Cordi et al., 2020). Thus, by initiating mental concepts related to “stress” or “sleep”, these concepts remain active during ongoing sleep and either impair or improve neurophysiological sleep regulation, respectively. Generally, the theoretical framework of “embodied cognition” offers an explanation for mutual influences between cognitive processes and body-related functions (Shapiro, 2010). According to this notion, mental processes involve simulations of body-related perceptions and actions, and therefore a close mutual link exits between cognitive processes and body-related sensorimotor systems of the brain (Gallese and Lakoff, 2005; Goldman, 1992). Evidence for embodied cognition is provided, for example, by brain imaging studies showing that concrete and abstract verbal concepts are capable of activating concept-related sensory, limbic and sensorimotor areas in the brain: e.g. the words “arm” vs. “leg” activate the respective sensorimotor area (Boulenger et al., 2012). Conversely, a higher compatibility between processed and performed action (action-compatibility effect; Glenberg and Kaschak, 2002; Zwaan et al., 2012) or a pre-activation of motor areas (via movement priming or neurostimulation) facilitates processing of action words in classical language comprehension areas of the brain (e. g. Mollo et al., 2016; Pulvermüller et al., 2005). While there is abundant evidence supporting the embodiment of action-related concepts, the embodiment of sleep-related concepts has not yet been examined. Evidence for embodied processing of sleep-related concepts would provide an important aspect to take into consideration for possible mechanisms underlying sleep-promoting effects of verbal and suggestive relaxation techniques.

Here we conducted a first study examining whether the processing of sleep-related concepts is “embodied”. Similar to the “action-compatibility effect”, we examined whether the “sleep-compatibility” of our body posture would influence the processing of sleep-related words. As sleep occurs almost exclusively in a relaxed lying body position, we expected a higher “sleep compatibility” for the processing of sleep-related concepts when participants were lying down and relaxing compared with an activated and upright position. In our study, participants listened to sleep- or activity-related words at four different volume levels close to their speech recognition threshold and had to categorise these words (see Figure 1a, for a summary of our procedure). To assess how difficult it was to process the meaning of the word, we focussed on event-related potentials (ERPs), with the N400 as our primary outcome parameter. The N400 amplitude in response to a stimulus is generally larger when the content of the stimulus is unexpected in a given context, thereby representing the ease of lexical access to the meaning of a word (Kutas and Federmeier, 2011; Lau et al., 2008). Therefore, we hypothesised that the N400 will be smaller when processing of word categories is compatible with the body position (congruent condition: sleep words – lying down; activity words – standing upright) compared with an incompatible body position (incongruent condition: sleep words – standing upright; activity words – lying down). In addition, we hypothesised that on the behavioural level, categorisation performance of the words would be better in the congruent compared with the incongruent condition. We expected the embodiment effects, both on the electrophysiological level as well as on a behavioural level, to generally be rather small and to primarily occur in processing conditions with a very high difficulty, i.e. at the lowest volume levels of the presented words.

**Figure 1:**
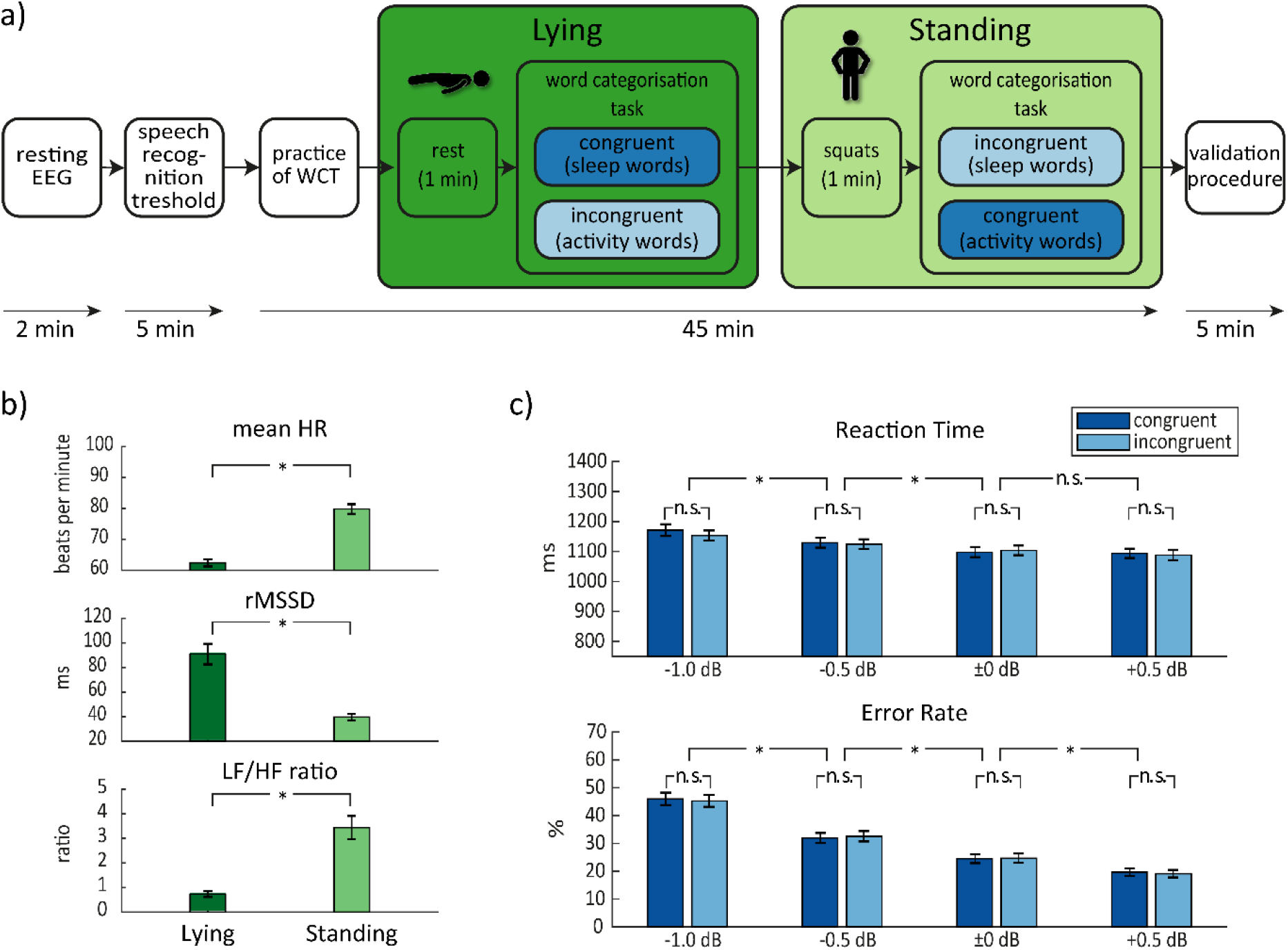
(a) Sequence of the experimental procedure. Subjects were seated in a sound attenuated and electrically shielded recording cabin. First, a two-minute resting EEG was recorded. Then the individual speech recognition threshold was acquired while participants were sitting. Participants practiced the word categorisation task (WCT) while still sitting. They were then asked to lie down on a bed in the recording cabin and to breathe calmly and deeply for one minute. The first block of the word categorisation task followed while participants were lying. Participants where then asked to stand up and do squats for one minute. The second block of the word categorisation task followed while participants were standing. The order of the positions (lying vs. standing) was balanced across participants. Subsequently, in a validation procedure, stimulus material was presented visually and rated regarding category, valence, arousal and imaginability. This was done while sitting. The cognitive tasks took roughly one hour, while the whole procedure including preparation and debriefing took two and half hours. (b) Effect of body position (lying vs. standing) on heart rate (HR) and heart rate variability (HRV) during the word categorisation task. Mean HR (beats per minute, top) was reduced and rMSSD (root mean square of successive differences, middle) was increased during lying compared with standing. LF/HF ratio decreased during lying compared with standing, possibly indicating relatively more parasympathetic influence compared to sympathetic influences in the lying compared with the standing condition. (c) Mean reaction times and error rates according to stimulus onset in the word categorisation task. In contrast to our hypotheses, congruency of the body position neither affected reaction times nor error rates. Generally, categorisation performance was better for words presented at a higher compared with lower volumes, both for reaction times and error rates. Means ± SE are indicated. *: p < .05 for planned (b) and exploratory (c) pairwise comparisons.

## 2. Material and methods

### 2.1 Participants

The final sample size of 66 (50 female, 16 male) participants was comprised of both students from Fribourg university as well as young professionals. Mean age ± standard deviation (SD) was 23 ± 4 years and ranged from 19 to 40 years. Initially 71 subjects were invited to participate in the study. Four subjects were excluded due to a technical problem by which no responses were registered during the main task. One subject had to be excluded due to excessive sweating artefacts in the electroencephalogram (EEG).

Preconditions for eligibility were the following: no hearing damage (e. g. tinnitus), age between 18 and 40 years old, right handedness, no acute health problems, no neurological or psychological impairments (e. g. depressive mood), no surgery within the past 3 months, no regular intake of medication (e. g. insulin therapy for diabetes) except for oral contraception, no acute sleep disorders, at most moderate alcohol consumption according to world health organisation (WHO) guidelines, at most moderate tobacco use on maximally four days a week, and at most minor cannabis use on maximally three days a week. Women were neither pregnant nor breast-feeding.

Participants guaranteed completeness and accuracy of the information they provided and gave written informed consent prior to participation. The study was approved by the local ethics committee and is in accordance with the Declaration of Helsinki (World Medical Association, 2001). Participants received either course credits or 30 Swiss francs for participation.

### 2.2 Experimental procedure

The study took place in the sleep laboratory of the University of Fribourg. Subjects arrived either in the morning between 08h45 and 09h15 or in the afternoon between 16h30 and 17h15. Their eligibility to participate in the study was determined by questionnaires. Upon arrival, experimental procedures and information about the alleged aim of the study were provided. Participants were told that we wanted to examine the recognition performance of acoustic stimuli close to the speech recognition threshold and furthermore, that we wanted to examine whether body position had an additional influence on their recognition performance. However, the concept of embodiment was not mentioned. In particular, it was not mentioned that the auditory stimuli were related to the body positions.

Participants filled out trait questionnaires regarding their chronotype (Griefahn et al., 2001), sleep quality (PSQI, Buysse et al., 1989), extraversion (NEO-FFI extraversion scale, Borkenau and Ostendorf, 1993), stimulus reducing-augmenting (Vando RA-scale, Barnes, 1985), and absorption (TAS, Ritz and Dahme, 1995). These are of no further relevance for the present analyses.

Subsequently they were seated in a sound attenuated and electrically shielded recording cabin and were prepared for simultaneous EEG, electrooculogram (EOG), and electrocardiogram (ECG) recordings. The experimenter, to guarantee a firm hold, attached ear clip headphones to both of the participant’s ears.

Participants filled out a mood questionnaire (Steyer et al., 1997) before starting with the cognitive tasks. As this mood questionnaire contained a sleep-related word that was used later on in the main task, a second mood questionnaire was constructed with the same amount of activity-related words, used in the main task. For the cognitive tasks, all instructions were given via a computer screen or via the headphones. However, the experimenter could be contacted at any time, in case of further questions. First, a two-minute resting EEG was recorded (one minute eyes open and one minute eyes closed in a randomised order across participants). Then the individual speech recognition threshold was acquired with a staircase procedure while participants were sitting (for details on the staircase procedure see supplementary material).

Participants practiced the word categorisation task while still sitting. Then subjects were informed that they had to execute the word categorisation task in two positions, lying versus standing (Figure 1a). Order of positions was counterbalanced across subjects. They were informed that the difficulty of the task would be that some of the words would be presented very quietly. All following instructions were then given via the headphones. First subjects were instructed to place themselves into the first position (lying down on a bed or standing upright), then subjects were informed that they should either breathe calmly and deeply (position lying) or do squats (position standing) for one minute. Subjects could start the countdown of the minute individually and were informed when a minute had passed. Afterwards the word categorisation task started. Subsequently, in a validation procedure, the words used in the word categorisation task were presented visually and participants rated them according to category, valence, arousal and imaginability without any time restriction (see supplementary material for results of the validation procedure). The validation was done while sitting. Finally, participants filled out the original mood questionnaire for a second time and were then debriefed, in which they were informed about the embodiment hypothesis. The cognitive tasks took roughly one hour, while the whole procedure took two and half hours.

### 2.3 Cognitive tasks

Cognitive tasks (speech recognition threshold, word categorisation task, validation procedure) were presented on a 19” LCD monitor (HP Compaq LA 1956x) with 1280 x 1024 resolution and a 60 Hz refresh rate using E-Prime presentation software (Eprime 2.0, Psychological Software Tools, Pittsburgh, PA). The computer volume was set to 50 % and the ear clip headphones (Philips SHS4700) were connected to the computer via an extension cable.

### 2.4 Word categorisation task

Participants heard spoken words belonging either to the category sleep or to the category activity. They had to decide which category the presented word belonged to. Subjects were informed that in some trials only noise would be presented. If they had heard noise or had not understood the word, they should not respond. The stimuli were presented at E-Prime volumes −500, −250, ±0, and +250 relative to the individual speech recognition threshold. These values correspond to roughly −1.0 dB SPL, −0.5 dB SPL, ±0 dB SPL, and +0.5 dB SPL relative to the individual speech threshold.

The task consisted of one minute of deep breathing, 12 minutes categorising while lying down, one minute of doing squats, and 12 minutes of categorising while standing upright. A total of 480 trials were presented (240 while lying and 240 while standing). Nine unique sleep-related words, nine unique activity-related words (see Table 1), and two unique noise files (Brown Noise) were presented at four different volume levels (= 80 stimuli). The presentation of the 80 stimuli was repeated three times (=240 stimuli). Stimuli were presented in random order within each of the three repetitions.

**Table 1:** List of original, German stimulus material in the word categorisation task and its translation to English. Words are alphabetically ordered. *: The words “groggily” and “sporty” were also presented in the mood questionnaires that were filled out by the participants before starting with the cognitive tasks.

| sleep-related words |  | activity-related words |  |
| --- | --- | --- | --- |
| dösen | *to doze* | anfeuern | *to cheer so. on* |
| Kissen | *pillow* | Eile | *rush* |
| schlapp* | *groggily* | Hochleistung | *high performance* |
| schnarchen | *to snore* | hüpfen | *to jump* |
| Siesta | *siesta* | rumtoben | *to romp* |
| Trägheit | *inertness* | Schnelligkeit | *speed* |
| träumen | *to dream* | sportlich* | *sporty* |
| liegen | *to lie* | stehen | *to stand* |
| Tiefschlaf | *deep sleep* | Stress | *stress* |

Each trial began with showing a fixation cross on the centre of the screen for a variable time ranging between 100 and 500 ms. Then the audio file was presented. Each unique word stimulus had an onset delay of 101 to 132 ms; Brown noise had an onset delay of 30 ms. After the audio onset the fixation cross changed into a question mark. Subjects had a maximum of 2400 ms to respond, starting from ca. 100 ms after stimulus onset. After the participants’ response a fixation cross was shown for 2400 ms minus the reaction time. Therefore, subjects could not decrease the duration of the task by answering especially fast. The computer screen was not needed for executing the task as all instructions and relevant stimuli were presented via the headphones. However, participants had the possibility to see the computer screen when executing the task while standing, but could not see the screen while lying.

A number pad was used as response device (Gembird USB NumberPad KPD-01). Participants were instructed to hold the number pad in their non-dominant hand and respond with the index finger of their dominant hand (all participants were right-handed). The left key “4” was always mapped to the first category and the right key “6” was always mapped to the last category. First and last relates to the order of mentioning the categories in the instructions. This order was counterbalanced across subjects. They were briefed to rest their index finger in between both response keys in between trials to guarantee accurate reaction time calculation. An animated figure was shown to visualize the response keys and accurate behaviour. Subjects were instructed to respond as fast and as accurate as possible.

The stimuli were generated with a text-to-speech software provided by Microsoft Cognitive Services (https://microsoft.com/cognitive-services/text-to-speech/#features, retrieved 2016/2017), using a male German (Austria) voice. Words were adjusted to have a 750 ms duration. More information on the selection and validation of the stimulus material can be found in the supplementary material.

Participants first practiced the task while sitting. The practice contained a minimum of 12 and a maximum of 84 trials. It ended as soon as subjects responded in 80 % of the ongoing trials correctly. During practice, participants received feedback on the correctness of their responses.

### 2.5 Manipulation check: effects of body position on HR and HRV

ECG was continuously recorded with silver-silver chloride (*Ag/AgCl*) electrodes (55 mm diameter, Kendall H98SG, Covidien). Two bipolar ECG channels were placed according to the standard lead II configuration. ECG was amplified by a BrainAmp ExG amplifier with an input impedance of 10 MΩ (Brain Products GmbH, Munich, Germany). Recordings, in AC mode, were sampled at 1000 Hz with a 10 μV resolution. The pass-band of the hardware filter was set to .016 to 250 Hz (see next section for further details).

Heart rate (HR) and heart rate variability (HRV) were analysed using the software Kubios HRV Premium Version 3.2.0 (Tarvainen et al., 2014). Segments for both positions, consisting of 12 minutes continuous data each (recorded during the execution of the word categorisation task) were processed separately. We used the built-in QRS detection algorithm and the automatic correction procedure to correct for artefacts and ectopic beats (see Kubios HRV User’s Guide, section 3.2, available at http://kubios.uef.fi for details). This automatic procedure lead to a median of 0.55 % detected artefacts (range: 0.30-11.65 %). We exported mean HR (in beats per minute), root mean square of successive differences (rMSSD in ms), as well as the ratio between the power of the low frequency (LF) component (0.04-0.15 Hz) and the power of the high frequency (HF) component (0.15-0.40 Hz) of HRV (LF/HF ratio). Power values were calculated using the fast Fourier transform (FFT) procedure.

### 2.6 EEG recording and quantification

EEG was continuously recorded with an Acti-Cap electrode system (EasyCap GmbH, Herrsching, Germany) from 64 sites positioned according to the 10-10 electrode reference system (Chatrian et al., 1985). All sites of EEG were referenced online to FCz. AFz served as ground. Silver-silver chloride *(Ag/AgCl)* electrodes were used. EEG was amplified by a BrainAmp DC amplifier with an input impedance of 10 MΩ (Brain Products GmbH, Munich, Germany). Recordings, in AC mode, were sampled at 1000 Hz with a 0.1 μV resolution. Impedances of EEG electrodes were kept below 15 kΩ. The pass-band of the hardware filter was set to .016 to 250 Hz (high pass filter: first-order filter with – 6 dB/octave; low pass filter: fifth-order Butterworth filter with −30 dB/octave). Recorded data was stored on a hard disk for later processing. EEG data was processed in MATLAB (R2018b, The MathWorks, Inc.), using the EEGLAB toolbox (v2019.1, Delorme and Makeig, 2004).

Raw EEG data was first pre-processed for running independent component analysis (ICA). ICA was used for removing eye, muscle and heart artefacts. Data was high pass filtered (1 Hz) and low pass filtered (30 Hz) using a zero-phase Hamming-windowed sinc FIR filter implemented in EEGLAB (eegfiltnew.m; Widmann et al., 2015). Then the two blocks of the word categorisation task were segmented and concatenated, excluding the one minute breathing and one minute squats. Noisy channels were rejected using the artefact subspace reconstruction (ASR) method (Chang et al., 2018; Kothe and Jung, 2014). Channels were rejected if they had flat line periods exceeding 5 seconds, were correlated less than .8 to their neighbouring channels, or had line noise exceeding 4 SD. On average 3.74 (median: 2) channels were removed in each data set ranging from 0 to 18 removed channels (mainly Fp and AF channels, which were therefore not used for later analyses). EEG data was re-referenced to the average reference after noisy channel removal and the online reference channel FCz was re-used in the data. Cz was discarded after re-referencing to match the number of channels and the rank of data for ICA. Cz was chosen because its activity is best captured by neighbouring electrodes and presumably the least amount of information is lost by discarding this electrode. Data was temporally segmented into consecutive one-second epochs and artefactual epochs were rejected automatically using joint probability and kurtosis (Delorme et al., 2007). Rejection criteria for joint probability and kurtosis were set to 5 SD for single channels and 2 SD for all channels. Rejection of an epoch always led to the rejection of this epoch in all channels. This automatic procedure lead to an average of 14 ± 2 % of rejected epochs (range: 9-18 %). ICA was performed on the remaining segments, using the AMICA algorithm (Delorme et al., 2012). The resulting weight matrix was saved.

In a second step, raw EEG data was pre-processed for analysing event-related potentials. Raw EEG data was high pass filtered (0.1 Hz) and low pass filtered (30 Hz) using a zero-phase Hamming-windowed sinc FIR filter implemented in EEGLAB (Widmann et al., 2015). Then the two blocks of the word categorisation task were segmented and concatenated, excluding the one minute breathing and one minute squats. Noisy channels, detected by the previous ASR procedure (described above) were removed, data was re-referenced to the average reference, the online reference channel FCz was re-used in the data, and Cz was discarded. The ICA weight matrix was imported and ICs were automatically classified using ICLabel (Pion-Tonachini et al., 2019). ICLabel provides the probability of each component to be generated by brain, muscle, eye, heart, line noise, channel noise, and other. We removed the ICs with highest probability of being generated by the eye (if probability exceeded 90%), the muscle (if probability exceeded 25%), and the heart (if probability exceeded 60%). Artefactual ICs were removed by setting their weights to 0 and re-calculating EEG activity with the resulting weight matrix. Thereafter rejected channels were interpolated (spherical). Trials were extracted by segmenting −200 ms to +1800 ms around stimulus onset. Each trial was baseline corrected by subtracting the mean baseline activity (−200 to 0 ms according to stimulus onset) from all data points of the trial. Finally, artefactual epochs were rejected automatically using the same method and parameters as described above. This automatic procedure lead to an average of 17 ± 3 % rejected epochs (range: 7-23 %). Only trials with correct responses were kept for the event-related potential analysis. Furthermore, the first 4 presentations of each unique word stimuli were discarded (containing each volume level once), as these first presentations showed pronounced different brain response compared to the stimuli that followed. Electrode positions Fp1, Fp2, AF7, AF3, AFz, AF4, AF8, O1, Oz, O2, and Iz were not included in the analyses because they were often noisy, and interpolation might have been be poor due to their location.

Separate averages were computed for each individual, electrode, volume level (−1.0 dB, −0.5 dB, ± 0.0 dB, + 0.5 dB), as well as for congruent (sleep words while lying and activity words while standing) and incongruent trials (sleep words while standing and activity words while lying). Amount of trials in the congruent vs. incongruent conditions were equalised individually for each subject and each volume level by choosing *n* random trials from the condition with the higher amount of trials *(n* being the amount of trials in the condition with lower amount of trials).

By visually inspecting the time course of the topography averaged over all participants and conditions, we detected three components in response to the speech stimuli. The exact temporal extent and maximum of each component was defined by inspecting the grand average event-related potential waveform. Suitability of the chosen time windows were checked and subsequently confirmed for each participant individually. Mean amplitude was calculated for each of the three components in the corresponding time windows (Luck, 2014), for each individual, electrode, volume level (−1.0 dB, −0.5 dB, ± 0.0 dB, + 0.5 dB), as well as for congruent and incongruent trials.

### 2.7 Statistical analyses

All statistical analyses were conducted with IBM Statistics for Windows version 26 (SPSS, Inc., IBM company) or MATLAB (R2018b, The MathWorks, Inc.), except otherwise specified. Significance levels were set to p_two-tailed_ < .05. Violations of sphericity were appropriately corrected by Greenhouse-Geisser ε_GG_ if ε_GG_ ≤ .75 (Box, 1954; Geisser and Greenhouse, 1958) or Huynh-Feldt ε_HF_ if ε_GG_ > .75 (Huynh and Feldt, 1976). Post hoc t-tests were conducted for further analyses of significant results. The measure of effect size ω^2^ is reported for significant results (Hays, 1994). It is an estimator for the population effect Ω2, which specifies the systematic portion of variance in relation to the overall variance (Rasch et al., 2014a, 2014b). All power calculations were done with G*Power (version 3.1.9.4, Faul et al., 2007) or MATLAB (R2018b). The sample was balanced according to starting time of the experimental session, order of positions, mapping of response keys, and sex. These factors were therefore controlled for and not included in statistical analyses. Data is always presented as mean ± standard error (SE) except otherwise specified.

#### 2.7.1 Manipulation check: effects of position on HR and HRV

Three dependent sample t-tests were used to compare cardiovascular measures (heart rate, rMSSD, and LF/HF ratio) in the lying versus standing condition.

#### 2.7.2 Word categorisation task – behavioural data

Behavioural results (dependent variables: reaction time, error rate) in the word categorisation task were analysed by a 2 x 4 ANOVA with the within-subjects factors condition (congruent vs. incongruent) and volume (−1.0 dB SPL, −0.5 dB SPL, ± 0 dB SPL, +0.5 dB SPL relative to the individual speech recognition threshold) in a repeated measure design. For reaction time analysis only trials with correct responses were included. Furthermore, reaction time distributions for each subject were calculated. Reaction times which exceeded the criterion of three interquartile ranges above the third quartile of the individual distribution (Tukey, 1977) or that were faster than 200 ms, were deemed outliers and removed for statistical analyses.

#### 2.7.3 Word categorisation task – event-related potential data

We conducted mass univariate analyses (Groppe et al., 2011) separately for the three detected ERP components (N180: 130-230 ms, P280: 230-330 ms, N400: 400-1800 ms in fourteen 100 ms sub-epochs). Separate repeated measure ANOVAs were calculated for each electrode (54 electrodes included) and each 100 ms sub-epoch, including the factors condition (congruent vs. incongruent) and volume (−1.0 dB SPL, −0.5 dB SPL, ± 0 dB SPL, +0.5 dB SPL relative to the individual speech recognition threshold). In case of significant effects, we conducted post hoc t-tests for the congruency by volume interaction (i.e. testing the congruency effect separately for each volume level) and the volume main effect. We corrected for multiple comparisons using cluster-based permutation tests (Groppe et al., 2011; Maris and Oostenveld, 2007). A cluster consists of electrodes connected in time and space that show significant effects. Only electrodes showing effects in the same direction were included in a cluster. The test reveals whether a cluster of a specific size is likely to be present in data where the condition labels of trials have been randomly shuffled. If the size of a cluster found in the original data is highly unlikely in permuted data, the cluster is classified significant. Importantly, cluster-based permutation tests neither reveal the exact time period when significant differences are present nor the exact location of the detected effect (Sassenhagen and Draschkow, 2019). We included 10’000 permutations and used F-mass/t-mass (sum of all F-/t-values taken from electrodes showing significant effects (inferred from parametric statistics described above)) and cluster size (amount of significant electrodes for each time period) to quantify cluster significance.

## 3. Results

### 3.1 Manipulation check: effects of position on HR and HRV

We first tested whether our manipulation of body position (lying down vs. standing upright) during the performance of the word-categorisation task resulted in the expected differences in arousal markers. Consequently, HR and HRV measures were derived from the twelve minute segments, while participants executed the word categorisation task. As expected in the lying position, subjects showed lower mean HR (beats per minute; t(65) = −14.62, p < .01, ω^2^ = .62), higher rMSSD (ms; t(65) = 6.96, p < .01, ω^2^ = .26), and lower LF/HF ratio (t(65) = −2.71, p < .01, ω^2^ = .21) compared to standing (Figure 1b). Manipulation of the body position therefore led to the expected differences in HR and HRV in the lying compared with the standing position. Thus, our manipulation of body position was successful and resulted in a calm vs. activated body state for lying down and standing upright respectively.

### 3.2 Behavioural data

In the next step, we analysed whether the difference in body positions affected performance in the detection success of the word-categorisation task. According to our hypothesis of the embodiment of sleep- and activity-related words, we expected the categorisation of sleep-related words to be better during a calm body state (i.e. lying down). In contrast, the categorisation of activity-related words should profit from an activated body state (i.e., standing up). Thus, categorisation performance should be generally better for words congruent to the body position (i.e., sleep-related words whilst lying down and activity-related words whilst standing up). In contrast, categorisation should be generally worse for words incongruent to the body position. We expected the effect of congruency to be present in highly difficult perception conditions, i.e. in our study predominantly when the volume was below the speech recognition threshold. Therefore, our main primary outcome was the interaction between the factor congruency (congruent vs. incongruent) and the factor volume (−1.0 dB SPL, −0.5 dB SPL, ±0.0 dB SPL, +0.5 dB SPL).

In contrast to our hypothesis, we did not find a significant interaction between the factor congruency and volume, neither for reaction times (F(1,65) = 1.37, p = .26) nor error rates (F(3,195) = .78, p = .51). Furthermore, we also did not find a significant main effect of congruency (reaction times: F(1,65) = 1.37, p = .25, error rates: F(3,195) = .08, p = .77). The ANOVA had a minimum of 95 % power to find effect sizes of Ω^2^ ≥ .01 for the main effect congruency and the interaction congruency by volume, both for reaction times and error rates. We can safely exclude the existence of a small and larger effect of congruency between body position and word category on the categorisation response to sleep- and activity-related words. Thus, the congruency between body position and word category did not influence the categorisation speed nor categorisation success, and this was not modulated by the volume of the presented words.

As expected, categorisation performance was better for words presented at a higher volume compared with a lower volume. The effect was present for both reaction times (F(3, 195) = 42.03, p < .01, ω^2^ = .32) and error rates (F(3, 195) = 305.06, p < .01, ω^2^ = .78). All volume levels differed significantly from each other with the exception of ±0.0 dB and +0.5 dB with regard to reaction times (see Figure 1c, for post hoc tests).

### 3.3 Event-related potential (ERP) data

#### 3.3.1 Characterisation of the general event-related response to aurally presented words

We detected three components in response to the speech stimuli, which were presented at volumes around the individual speech recognition threshold (Figure 2). Firstly, a centro-parietally distributed negative wave, ranging from 130 to 230 ms and peaking at 180 ms (N180). Secondly, a bilateral parieto-temporally distributed positive wave, ranging from 230 to 330 ms and peaking at 280 ms (P280). N180 and P280 were strongly and globally influenced by volume. The mean amplitude of both components increased with stimulus intensity (Figure 2b), as is typical for early sensory ERP components reflecting the physical properties of the stimuli (e. g. volume). Thirdly, we detected a frontally distributed, slightly left oriented N400 peaking at 800 ms. This component emerged at 400 ms and slowly faded out. The N400 was reflected as a positively oriented wave in the parieto-occipital cortex. Moreover, we detected two distinct volume effects on the N400 (see Figure S5 in the supplementary material). Initially, the N400 was less pronounced for the lowest volume level compared with all other volume levels. Latterly, in contrast to the early exogenous components, the mean amplitude of the endogenous N400 component decreased with stimulus intensity: lower volumes led to a higher mean activity of the N400 compared with higher volumes. This indicates that it is more difficult, and it takes longer, to infer the meaning of a word presented at volumes below the speech recognition threshold. Descriptively the lowest volume always had the lowest mean activity for the N180 and P280, with increasing mean amplitude as volume increased. In contrast, for the N400, mean amplitude was always largest for the lowest volume and continuously decreased with increasing volume. Post hoc tests for the volume effect revealed that the mean amplitude of the N180 and P280 differed from each other for all volume levels, while for the N400 mean amplitude mainly differed for the lowest volume level (−1.0 dB SPL) compared with all other volume levels. Detailed analyses of the volume effect can be found in the supplementary material.

**Figure 2:**
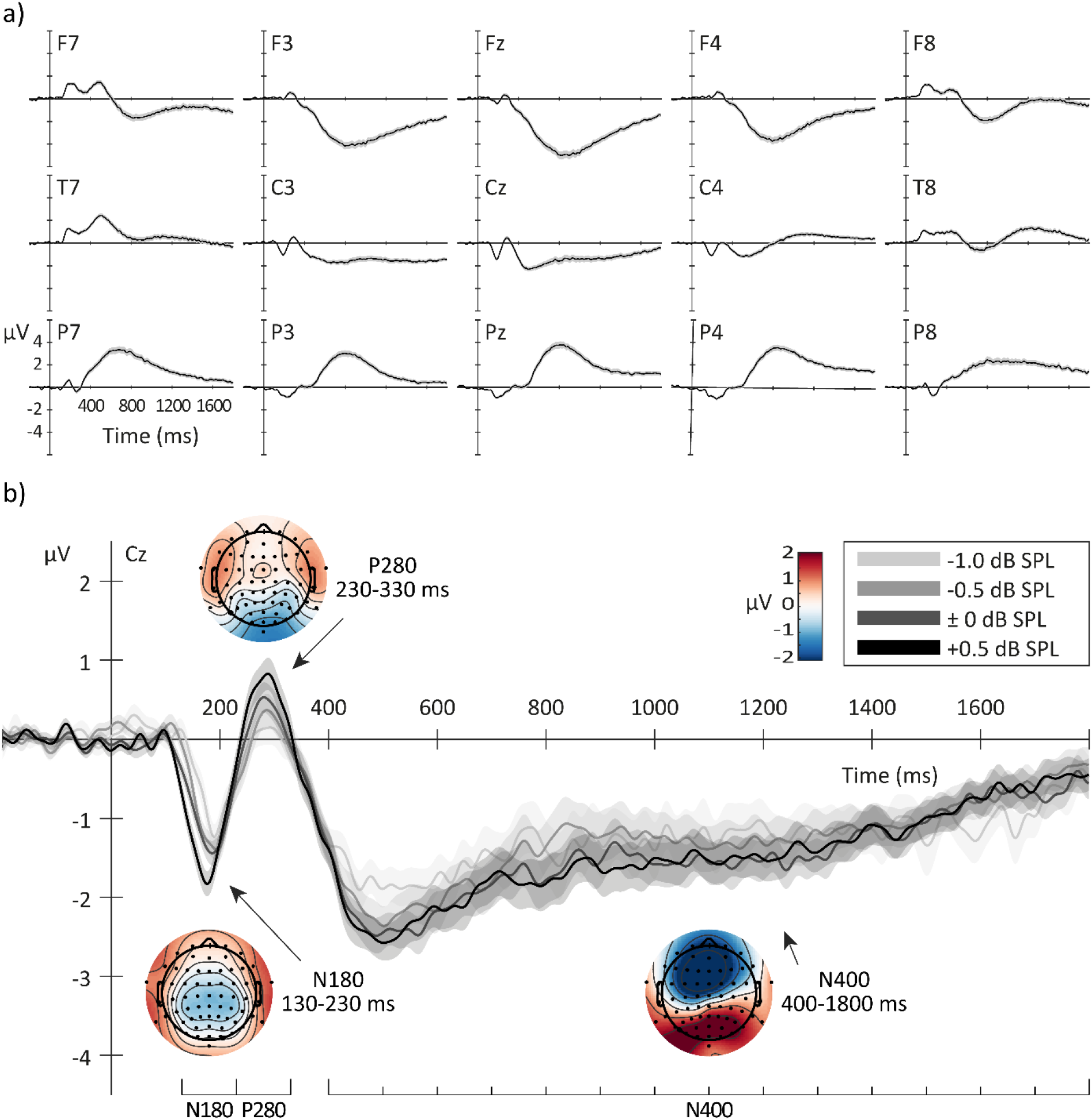
(a) Grand averages of the ERP (± SE) waveforms for a representative selection of electrodes averaged over all conditions and participants. The amplitude of N180 and P280 are relatively small compared to the N400 component which can be clearly detected as a negative going wave in frontal and central positions (upper two rows) and as a positive going wave in the parietal cortex (lower row). The N180 and P280 can be best detected in the central electrodes close to the midline. (b) Grand average of the ERP (± SE) at Cz, averaged over all participants and congruency conditions separated for the four different volume levels. The mean amplitude of the N180 and P280 increases with stimulus intensity (light grey – lowest volume to dark grey – highest volume). See supplementary material for details on the volume effects. Additionally, the mean topographic maps over all subjects, conditions, and time windows used for analyses of the three components emerging in response to the presented speech stimuli in the word categorisation task are depicted. The N180 is a centro-parietally distributed negative wave ranging from 130 to 230 ms and peaking at 180 ms. The P280 is a bilateral parieto-temporal distributed positive wave, ranging from 230 to 330 ms and peaking at 280 ms. The N400 (400-1800 ms) is a frontally distributed, slightly left oriented negativity, reflected as a positivity in the parietal cortex peaking at 800 ms (analysed in 14 consecutive 100 ms epochs). All timing information relates to the stimulus onset at 0 ms.

#### 3.3.2 Congruency effect on the N400

We examined whether congruency of body position and word category led to the predicted stronger negativity of the N400 for incongruent versus congruent trials. We expected the embodiment effect to be rather small and only to be present in highly difficult perception conditions. In our study this is related to the volume levels below the speech recognition threshold. Because the N400 spanned across a large time window, from 400 ms to 1800 ms after stimulus onset, and effects are not expected to be uniformly present over the complete period, we analysed mean activity in fourteen consecutive 100 ms epochs.

We found a significant cluster of electrodes, exhibiting a main effect of congruency for the N400 from 700 to 1100 ms (F-mass: 73.30, p < .05; cluster size: 13, p < .01). Mean amplitude of the N400 component was larger, that is more negative, for incongruent than for congruent trials. The contributing electrodes were located frontally close to the midline. The effect was maximal at FCz in the time window 700-800 ms (F(1,65) = 7.87, p < .01, ω^2^ = .05) and had the same direction for all significant electrodes and time windows. Effect sizes for the contributing electrodes ranged between .03 < ω^2^ < .05, with an average (± SD) of ω^2^ = .03 ± .01.

This main effect was further qualified by a congruency by volume interaction. A group of neighbouring electrodes over the right fronto-central cortex showed significant congruency by volume interactions. These electrodes formed a significant cluster from 400 to 800 ms (F-mass: 56.02, p < .01; cluster size: 16, p < .01) and a second significant cluster from 1200 to 1500 ms (F-mass: 35.61, p < .05; cluster size: 10, p < .05). For the early cluster (400-800 ms) the effect was maximal at F2 in the time window 700-800 ms (F(3,195) = 4.95, p = .01, ω^2^ = .02). Effect sizes for the contributing electrodes ranged between .01 < ω^2^ < .02, with an average (± SD) of ω^2^ = .01 ± .00. For the late cluster (1200-1500 ms) the effect was maximal at FC2 in the time window 1200-1300 ms (F(3,195) = 5.23, p = .01, ω^2^ = .02. Effect sizes for the contributing electrodes ranged between .01 < ω^2^ < .02, with an average (± SD) of ω^2^ = .01 ± .01.

Additionally, a parieto-occipital group of electrodes exhibited significant congruency by volume interactions. These electrodes formed a significant cluster from 500 ms to 1700 ms (F-mass: 130.80, p < .01; cluster size: 34, p < .01). The effect was maximal at P5 in the time window 900-1000 ms (F(3,195) = 6.66, p < .01, ω^2^ = .03). Effect sizes for the contributing electrodes ranged between .01 < ω^2^ < .03, with an average (± SD) of ω^2^ = .02 ± .01. However, this cluster should be interpreted cautiously, as it was mainly present due to a constant significant effect in the P5 electrode; none of the neighbouring electrodes contribute consistently to the cluster.

To further examine this interaction effect and its direction, we conducted post hoc t-tests. We tested the congruency effect separately for each volume level. Dependent sample t-tests indicated a main effect of congruency for the lowest volume level (−1.0 db SPL; Figure 3a). A group of neighbouring electrodes at the right-central frontal cortex showed a stronger negativity for incongruent than for congruent stimuli, forming a significant cluster (t-mass: 150.64, p < .05; cluster size: 56, p < 0.05). In this cluster of electrodes, the effect was present from 600 ms to 1600 ms. Mean amplitude of the N400 component was larger, that is more negative, for incongruent than for congruent trials. The effect was maximal at FC2 in the time window 700-800 ms (t(65) = 4.39, p < .01, ω^2^ = .12) and had the same direction for all significant electrodes and time windows (Figure 3b). Effect sizes for the contributing electrodes ranged between .02 < ω^2^ < .12, with an average (± SD) of ω^2^ = .05 ± .02.

**Figure 3:**
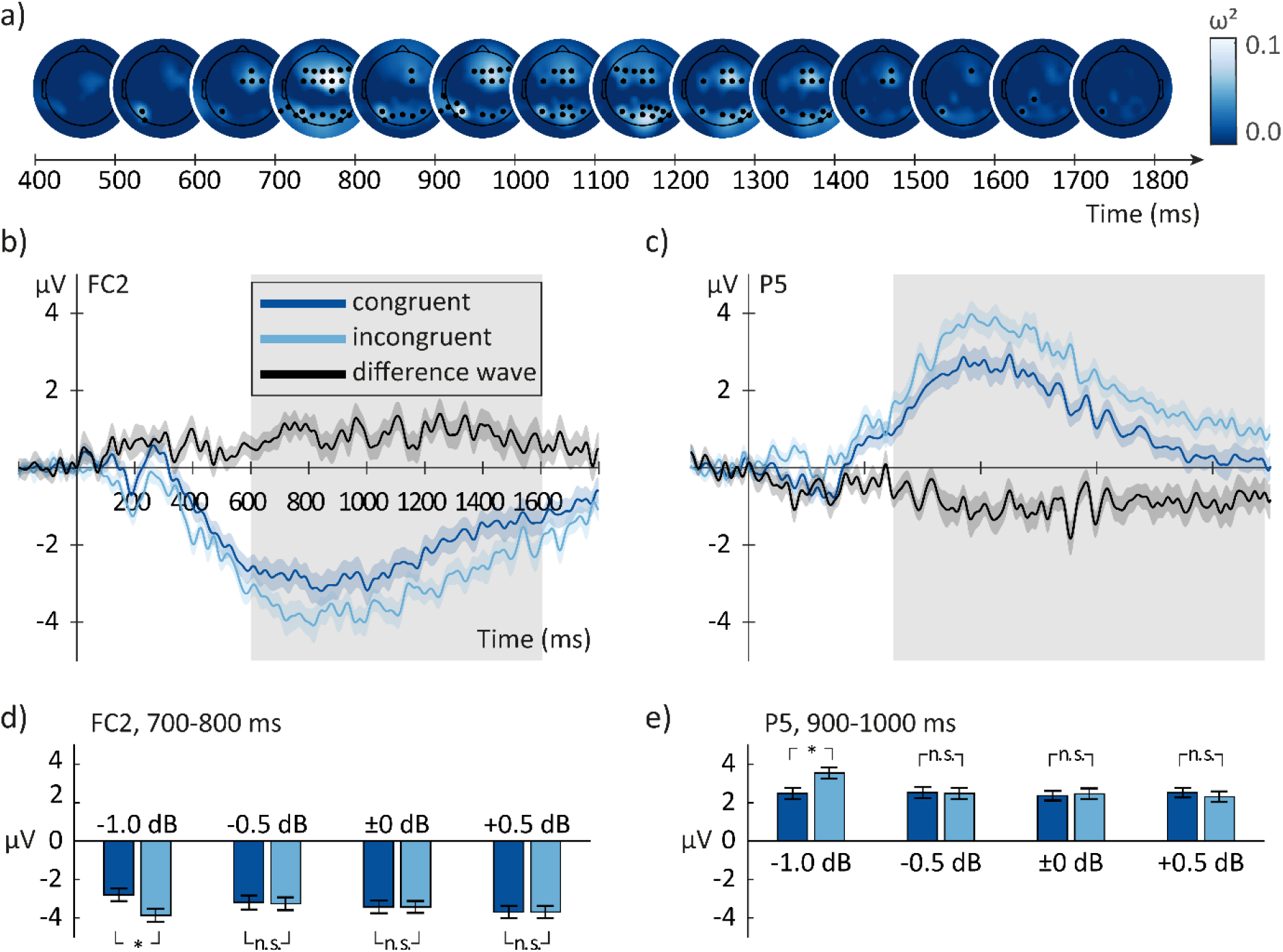
a) Topographic maps of the effect sizes (ω^2^) for the comparison of congruent versus incongruent trials at the lowest volume level (−1.0 dB SPL below the individual speech recognition threshold). Electrodes showing significant differences are plotted as black dots. The frontal cluster covers the right fronto-central cortex and emerges starting from 600 ms until 1600 ms. The parietal cluster is present from 500 to 1800 ms and spans both hemispheres. Lighter blue indicates larger effect sizes. Two exemplary electrodes from the frontal cluster (FC2, b)) and from the parietal cluster (P5, c)) showing the grand average ERP (± SE) for congruent (dark blue) and incongruent (light blue) trials, averaged over all participants for the lowest volume level (−1.0 dB SPL below the individual speech recognition threshold) are shown. Furthermore, the difference wave (± SE) between the two trial types is shown in black. The grey rectangle marks the period with significant differences between congruent and incongruent trials. The N400 amplitude is more pronounced for incongruent than for congruent trials, indicating that the processing of stimuli incongruent to the body position demands higher cognitive effort than the processing of stimuli congruent to the current body position. The bar plots show the mean N400 amplitude (± SE) for congruent (dark blue) and incongruent (light blue) trials averaged over the time window 700-800 ms for electrode FC2 (d)) and averaged over the time window 900-1000 ms for electrode P5 (e)). Congruent and incongruent trials differ in amplitude only within the lowest volume level. The chosen electrodes and the chosen time windows reflect the periods where the effect was maximal for the frontal and parietal cluster, respectively. All electrodes contributing to a cluster had the same effect pattern as depicted in these exemplary electrodes. *: p < .05.

As for the interaction effect of the ANOVA, the post hoc t-tests revealed an additional parieto-occipital cluster of electrodes exhibiting significant congruency effects from 500 ms to 1800 ms (t-mass: – 159.04, p < .05; cluster size: 62, p < .05; Figure 3a). The effect was maximal at P5 in the time window 900-1000 ms (t(65) = −4.45, p < .01, ω^2^ = .12; Figure 3c). Effect sizes for the contributing electrodes ranged between .02 < ω^2^ < .12, with an average (± SD) of ω^2^ = .04 ± .02. This cluster mirrors the frontal cluster. The N400 is reflected as a positively oriented wave in the parietal cortex. Here we observe a stronger positivity for incongruent compared with congruent trials. Therefore, as in the frontal cortex, incongruence leads to a larger mean amplitude of the N400. When analysing the lowest volume level alone, this cluster is less dependent on a single electrode (P5, cf. cluster in the interaction effect). Therefore, we will carefully interpret the parieto-occipital cluster. It supports the interpretation of the right-frontal cluster of significant electrodes and could be a reflection of the frontal effect.

Such a congruency effect could not be detected in any of the other three volume levels (−0.5 dB SPL, ±0 dB SPL, and +0.5 dB SPL; Figure 3d/e). Even though single electrodes showed significant congruency effects, none of these electrodes formed significant clusters and are therefore most likely chance findings. Across all t-tests, we achieved a minimum of 95 % power (calculated individually for each t-test and electrode) to find effect sizes (Ω^2^) of .05 ± .01 (range: .01 – .10) or larger. The ANOVA had a minimum of 95 % power (calculated individually for each electrode) to find effect sizes (Ω^2^) of .02 ± .01 (range: .00 – .05) or larger for the N400 main effect of congruency and .02 ± .01 (range: .01 – .04) or larger for the N400 congruency by volume interaction.

#### 3.3.3 Congruency effect on early components

To our knowledge there are no reports of congruency effects on early ERP components. We therefore did not have specific hypotheses of congruency effects on the first two ERP components.

Firstly, we examined the N180 component. We detected a significant cluster of electrodes at the parietal cortex showing a main effect of congruency for the N180 (F-mass: 58.49, p < .05; cluster size: 8, p < .05). These electrodes exhibited a stronger negativity for congruent than for incongruent stimuli. The effect was only present in the left hemisphere. The effect was maximal at P1 (F(1,65) = 10.57, p = .01, ω^2^ = .07) and had the same direction for all significant electrodes. Effect sizes for the contributing electrodes ranged between .03 < ω^2^ < .07, with an average (± SD) of ω^2^ = .05 ± .01.

Furthermore, this group of neighbouring electrodes showed a significant congruency by volume interaction, again forming a significant cluster (F-mass: 28.07, p < .05; cluster size: 7, p < .05). The effect was maximal at P5 (F(3,195) = 5.35, p < .05, ω^2^ = .02) and had the same direction for all significant electrodes. Effect sizes for the contributing electrodes ranged between .01 < ω^2^ < .02, with an average (± SD) of ω^2^ = .02 ± .00.

To further examine the N180 interaction effect and its direction, we conducted post hoc t-tests. We tested the congruency effect separately for each volume level. Dependent sample t-tests indicated a main effect of congruency for the lowest volume level (−1.0 dB SPL; Figure 4a). Similar to the effects of the ANOVA, a group of neighbouring electrodes on the left parietal cortex showed a stronger negativity for congruent than for incongruent stimuli, thereby forming a significant cluster (t-mass: −30.44, p < .01; cluster size: 10, p < .05). The effect was maximal at P1 and P3 (t(65) = −3.59, p < .01, ω^2^ = .08) and had the same direction for all significant electrodes (Figure 4b). Effect sizes for the contributing electrodes ranged between .02 < ω^2^ < .08, with an average (± SD) of ω^2^ = .06 ± .02.

**Figure 4:**
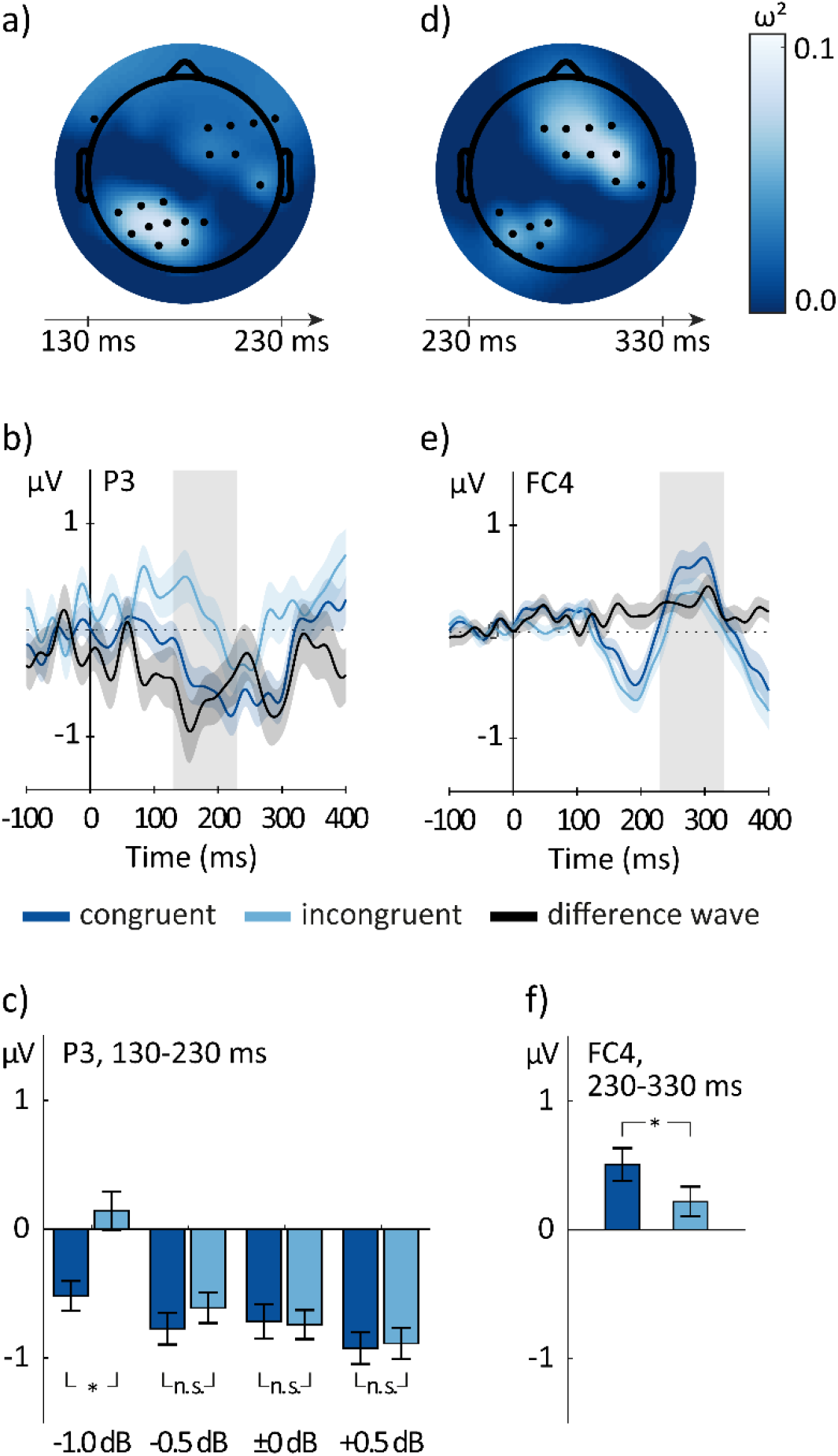
a) Topographic map of the effect sizes (ω^2^) for the comparison of congruent versus incongruent trials at the lowest volume level (−1.0 dB SPL below the individual speech recognition threshold) for the time window 130-230 ms post stimulus onset. Electrodes showing significant effects are plotted as black dots. A significant cluster was formed by a group of electrodes in the left parietal cortex. Lighter blue indicates larger effect sizes. b) Exemplary electrode (P3) from the parietal N180 cluster showing the congruency effect at the lowest volume level. The grand average ERP (± SE) for congruent (dark blue) and incongruent (light blue) trials is depicted as an average over all participants for the lowest volume level (−1.0 dB SPL below the individual speech recognition threshold). Furthermore, the difference wave (± SE) between the two trial types is shown in black. The N180 amplitude is more negative for congruent than for incongruent trials, indicating that speech stimuli congruent to the current body position are acoustically better understood than incongruent speech stimuli. The grey rectangle marks the period with significant differences between congruent and incongruent trials. c) Mean N180 amplitude (± SE) for congruent (dark blue) and incongruent (light blue) trials averaged over the time window 130-230 ms for electrode P3. Only within the lowest volume level do congruent and incongruent trials differ in amplitude. d) Topographic map of the effect sizes (ω^2^) for the main effect congruency of the conducted ANOVA in the time window 230-330 ms post stimulus onset. A significant cluster was formed by a group of electrodes in the right frontal cortex. e) Exemplary electrode (FC4) from the frontal P280 cluster showing the grand average ERP (± SE) for congruent (dark blue) and incongruent (light blue) trials, averaged over all participants and all volume levels. Furthermore, the difference wave (± SE) between the two trial types is shown in black. The P280 amplitude is more positive for congruent than for incongruent trials. Like for the N180, this could indicate that speech stimuli congruent to the current body position are acoustically better understood than incongruent speech stimuli. f) Mean P280 amplitude (± SE) for congruent (dark blue) and incongruent (light blue) trials averaged over the time window 230-330 ms and all volume levels for electrode FC4. The difference between congruent and incongruent trials is present in all volume levels. The chosen electrodes are those where the effect was maximal for the N180 and P280, respectively. All electrodes contributing to a cluster had the same effect pattern as depicted in these exemplary electrodes. *: p < .05.

The effect tended to be reflected in a marginally significant cluster in the right frontal cortex (t-mass: 15.55, p = .07; cluster size: 7, p < .05). Effect sizes for the contributing electrodes ranged between .02 < ω^2^ < .04, with an average (± SD) of ω^2^ = .03 ± .01. Here we observe a stronger positivity for congruent compared to incongruent trials. Therefore, as in the parietal cortex, incongruence leads to a larger mean amplitude of the N180.

Such a congruency effect could not be detected in any of the other volume levels (−0.5 dB SPL, ±0 dB SPL, and +0.5 dB SPL; Figure 4c). Across all t-tests, we achieved a minimum of 95 % power (calculated individually for each t-test and electrode) to find effect sizes (Ω^2^) of .06 ± .01 (range: .03 – .10) or larger. The ANOVA had a minimum of 95 % power (calculated individually for each electrode) to find effect sizes (Ω^2^) of .04 ± .01 (range: .03 – .05) or larger for the N180 main effect of congruency and of .03 ± .00 (range: .02 – .04) or larger for the N180 congruency by volume interaction.

Secondly, we examined the P280 component. A significant cluster of electrodes at the right frontal cortex showed a main effect of congruency for the P280 (F-mass: 80.42, p < .05; cluster size: 9, p < .05; Figure 4d). These electrodes exhibited a stronger positivity for congruent than for incongruent stimuli. The effect was maximal at FC4 (F(1,65) = 13.24, p = .01, ω^2^ = .08) and had the same direction for all significant electrodes (Figure 4e/f). Effect sizes for the contributing electrodes ranged between .02 < ω^2^ < .08, with an average (± SD) of ω^2^ = .06 ± .02. The effect was reflected as a negativity in the left parietal cortex, however these electrodes did not form a significant cluster (F-mass: 42.94, p = .06; cluster size: 7, p = .05).

For the P280, there was no congruency by volume interaction at any electrode. The ANOVA had a minimum of 95 % power (calculated individually for each electrode) to find effect sizes (Ω^2^) of .03 ± .01 (range: .02 – .04) or larger for the P280 main effect of congruency and of .02 ± .00 (range: .02 – .03) or larger for the P280 congruency by volume interaction. We can therefore exclude the possibility of the presence of small and larger effects.

### 4. Discussion

Here we present evidence from event-related potentials sustaining that the processing of sleep- and activity-related words is embodied. In particular, processing of words was facilitated in the congruent condition (sleep words – lying; activity words – standing) when words were presented at the lowest volume. This was indicated by a lower N400 amplitude in the congruent compared with the incongruent condition. As the N400 is typically associated with higher-order cognitive processing, we interpret the decrease in the N400 in the congruent condition as a facilitation of word processing caused by a compatibility of the body position under conditions where word processing is difficult. For earlier, more sensory ERP-components, the N180 and P280 showed larger deflections for congruent versus incongruent trials. Similarly, we observed larger deflections in these two early components with increased volume levels of the words. Thus, in addition to the facilitation of complex cognitive processing, a compatible body position might also improve sensory processing of words (e.g. the ease of understanding the word). We conclude that our ERP-results provide evidence for an embodiment of sleep- and activity-related words. In contrast to the electrophysiological level, we did not observe any effects of congruency between body position and word category on the behavioural level.

Our results of the embodiment of sleep-related words are in line with previous reports on the embodiment of action words. The action-compatibility effect predicts processing advantages, if processed and performed actions are compatible. In studies testing this effect, subjects are asked to judge the sensibility of sentences and respond with a movement. Response movements can be either congruent or incongruent to the movement described in the sentences. For example, Glenberg and Kaschak (2002) presented sentences implying a movement towards or away from the body. The response movement was significantly faster when its direction was compatible with the movement direction described in the sentence. Zwaan et al. (2012) presented sentences describing actions implying a forward or backward movement of the whole body. Forward-sentences caused a congruent shift of the body. For sentences implying a backward movement, the detected forward shift was much less pronounced compared to sentences describing a forward shift. This was found even though subjects had to respond by left vs. right movements to judge sensibility of the sentences.

Furthermore, movement priming or neurostimulation facilitate the processing of action words in classical language comprehension areas of the brain. In Mollo et al. (2016) participants either pressed a button with their index finger or a pedal with their big toe. The authors found that arm-related and leg-related words led to stronger activity in the posterior superior temporal gyrus and hand motor cortex when the word was congruent to the response type than when it was incongruent. Similar to our study, their manipulation led to rather small neurophysiological congruency effects which did not affect the behavioural responses. Pulvermüller et al. (2005) demonstrated that transcranial magnetic stimulation (TMS) applied to the arm- and leg-motor cortex facilitates the processing of congruent words (arm- and leg-related action words, respectively) when contrasted to incongruent words; i.e. responses to leg-related action words are faster compared with arm-related action words when the leg-motor cortex is stimulated 150 ms after visual word presentation.

These studies indicate that the pre-activation of motor networks facilitates the activation of compatible semantic networks. Conversely, semantic networks may be capable of activating corresponding motor networks. Evidence for the assumption of spreading network activation is for example provided by a study showing that listening to activity-related sentences can activate corresponding sensorimotor networks (Tettamanti et al., 2005). Two meta-analyses report that activated regions for action language processing and regions activated by motor execution, imagery, and observation are distinct, but overlap substantially (Courson and Tremblay, 2020; Yang and Shu, 2016). Therefore, these meta-analyses imply that language and action activate unique neuronal networks, which nevertheless overlap substantially and thereby are capable of activating each other.

In addition to movement priming, body positions have been successfully used as embodied “prime” in other studies, e.g. in the context of emotions (Price and Harmon-Jones, 2015). In particular, activity from facial muscles has proven to modulate our experiences of emotions. A recent meta-analysis found supporting evidence for the hypothesis that facial feedback influences emotions (Coles et al., 2019). Importantly, these influencing effects are rather small and heterogeneous in their nature. Evidence for a strong effect of facial feedback on emotions (Strack et al., 1988) could not be replicated in a multi-site replication project (Acosta et al., 2016), which fits well with our findings of rather small effect sizes and no effects of congruency on overt behaviour.

Pulvermüller and colleagues have proposed that words are represented in cortex-wide semantic networks which reflect concepts that are grounded in perception and action (Kiefer and Pulvermüller, 2012; Pulvermüller, 1995). Furthermore, Damasio’s somatic marker hypthesis assumes numerous interactions between body states and cognition (Damasio, 1994; Damasio, 1996). Similar to the studies on embodied language and emotion and based on these theoretical accounts we assume for our study that the body position of lying down induced a pre-activation of networks related to the semantic concepts of relaxation and sleep. Simultaneously, the processing of sleep-related words should activate the network representing the meaning of “sleep” but also bodily experiences, for example the experience of lying down and being relaxed. Subsequently corresponding words should be processed more efficiently than unrelated words.

The beneficial effect of a compatible body position on word processing occurred only at the lowest volume level for the N180 and the N400. The congruency effect for the P280 did not depend on volume. However, the differences between congruent and incongruent trials were descriptively larger for lower volumes compared to higher volumes. Generally, louder stimuli are easier to understand. Thereby, they quickly activate their corresponding semantic network. On the contrary, in highly difficult perception conditions, for example when playing the stimuli below the speech recognition threshold, the stimulus itself cannot quickly activate its corresponding semantic network. Therefore, if body position has pre-activated compatible semantic networks, this could help to process stimuli congruent to the current position. Consistent with this hypothesis, we mainly found embodiment effects at the lowest volume level. Furthermore, a compatible body position did not influence word processing on the behavioural level. Thus, the embodiment of semantic concepts might be of minor practical relevance for normal word processing during wakefulness.

Our result that a congruent body position facilitates processing of sleep words opens up the possibility that the influence could also act in the opposite direction: the activation of sleep-related words might in turn facilitate the activation of networks related to sleep induction and sleep maintenance. Sleep and wakefulness are controlled by mutual inhibitory influences between the wakefulness-promoting arousal-system and the sleep-promoting network (Saper et al., 2005). A multimodal representation of sleep might include the sleep-promoting network. Thus, as our results show that sleep-related words are embodied, an activation of these mental concepts might result in a more pronounced activation of the sleep-promoting network, thereby improving sleep quality. We assume in addition that the activity of mental concept can be initiated before sleep and remains active during sleep. An increased activity of a multimodal, embodied representation of the concept “sleep” during sleep could be a promising explanatory mechanism for the positive influence of pre-sleep cognitive intervention on later sleep processes (Cordi et al., 2014; Cordi et al., 2015; Cordi et al., 2020). Furthermore, we have recently provided evidence that experimental activation of the mental concept of “sleep”, by presenting relaxation-related words during sleep, is indeed capable of extending the time spent in slow wave sleep in young healthy participants (Beck et al., 2020). Thus, activation of multimodal representations of sleep-promoting networks is capable of influencing ongoing sleep processes. The study reported here adds to this notion by providing evidence that sleep-related concepts are in fact stored in a multimodal representation including body components, as body posture facilitates processing of sleep-related words.

Our study has several limitations. Firstly, we define the body position of “lying down” and a calm body state induced by deep breathing as congruent to sleep. We have seen that these are indeed key features associated with sleep (Saper et al., 2005). Nevertheless, our participants were fully awake and needed to be alert throughout the task. Not being actually asleep or active, but rather having induced a calm vs. activated body state could explain why we only found small congruency effects on ERPs as well as our findings specifically related to the lowest volume level. Future studies should test effects of embodiment of sleep-related concepts using manipulations more closely related to sleep, e.g. immediately waking up participants from the sleep state, comparing sleep deprived vs. non-sleep deprived subjects or examining potential differences in processing of sleep-vs. activity-related words during the sleep state itself.

Secondly, we successfully manipulated physical arousal, which was manifested in a lower HR and higher HRV for lying down versus standing upright positions. However, one might need to differentiate between physical and cognitive arousal (Blum et al., 1967). These two types of arousal could exert very different effects on sleep and cognition. For example, cognitive arousal has shown to impair sleep (Wuyts et al., 2012). On the contrary, physical arousal, in the form of exercise, does rather improve sleep (Kubitz et al., 1996; Stutz et al., 2019) and cognition (Chang et al., 2012). Thus, future studies might want the repeat the manipulation examining the effect of more or less cognitive arousal on the processing of sleep-related words.

Thirdly, the here elicited auditory N400 component deviates in latency and topography from the classic, visual N400 component. It is more frontally distributed (in contrast to the classic posterior topography) and rather late (peaking at 800 ms post stimulus) compared to the classical N400 literature. These differences are potentially caused by stimulus modality (auditory vs. visual) and type of incongruence: typically incongruence is studied as semantic incongruence within sentence settings. Here we have independent single words, presented auditorily, where incongruence is manipulated due to body position. However, the N400 is best characterised by a common functionality, but not so much by its localisation, time course or a specific mental operation (Kutas and Federmeier, 2011). Furthermore, literature suggests that the N400 component is not generated in a single structure but by various widely distributed cortical networks (Kutas and Federmeier, 2011; Lau et al., 2008) which could explain differences in topography depending on the study design.

Fourthly, the time-window of the N400 does overlap with the time-window where participants overtly responded with a button press. Therefore, we cannot exclude the possibility that the event-related potentials overlap with the movement-related cortical potentials. However, as each trial included in the analyses contains a button press, the differences in the event-related potentials are unlikely to be caused by participants’ responses. In addition, the N400 peak (800 ms post stimulus onset) considerably precedes the mean reaction time (1120 ± 16 ms post stimulus onset).

To conclude, our results support the prediction of the embodiment of sleep- and activity-related words. The effects of body position on mental processing of word meanings are small and do not affect behaviour measured in the form of reaction times and error rates. Future studies need to replicate our findings and further test the theoretical consequences of the embodiment of sleep-related words regarding current notions on the influence cognition has on sleep.

## Acknowledgements

We would like to thank Louisa Clarke for helpful comments on this manuscript. We furthermore thank our students for assistance in data collection and all subjects for their participation.

