## supplement for "Embodiment of sleep-related words: evidence from event-related potentials"

***Detection of the speech recognition threshold***

We identified the individual speech recognition threshold with a staircase procedure. Subjects had to decide whether they had understood the presented word or not in each trial. Subjects were informed that speed was of no relevance in this task. To highlight the importance of the difference between the hearing threshold and speech recognition threshold, we emphasised that having heard something was not relevant to the task, but that we were interested in whether participants had understood the presented word. The mean speech recognition threshold in the present sample is -5591 ± 338 E-Prime volume (computer volume set to 50 %).

A trial consisted of the presentation of a fixation cross for 250 ms. Subsequently a stimulus was presented for 750 ms while continuously showing the fixation cross for further 250 ms. Then a question mark was shown in the centre of the screen. Subjects could now respond by pressing a “yes”- or “no”-button on a number pad to indicate whether they had understood the presented word. The validity of participants’ responses was not controlled for. There was no time limit for responding.

The procedure always started with two words played at -1000 and -2000 E-Prime volume, which corresponds to clearly understandable stimuli. These two trials were not included into the threshold calculation, but served as familiarisation trials.

Then, three repetitions of the staircase procedure followed. The first trial of each repetition started with a randomly chosen volume of -2000 to -6000 E-Prime volume. This ranges from clearly to barely understandable. If a subject responded to have understood the first word, the following trials were played at a lower volume (-500). If a subject responded to not have understood the first word, the following trials were played at a higher volume (+500). In each trial the volume increased or decreased further. When volume was increased and the subject responded to have understood the word twice in a row, the following trials were played with a decreased volume (-500, direction change), and vice versa: when volume was decreased and the subject responded to having not understood the word twice in a row, the following trials were played with an increased volume (+500). The change of volume was adapted: first volume changed in steps of +/- 500, after two direction changes volume changed +/- 200, after further two direction changes volume changed +/- 100 and after further two direction changes volume changed +/- 50. In total one repetition consisted of eight direction changes (e.g. +500, -500, +200, -200, +100, -100, +50, -50) with a variable amount of trials for each subject between the direction changes (depending on the responses of the subject). The volume at direction change 5, 6, and 7 was averaged and saved. Finally, the saved values from all three repetitions were averaged to obtain the final individual threshold.

Fifty-four unique words, each adjusted to have a 750 ms duration were used in a balanced order. Each word was played one to four times, depending on the overall number of trials. Depending on the subjects' responses, the total amount of trials ranged between 93 and 167 trials (mean ± S. D.: 127 ± 15), not differing between the three repetitions (roughly 42 trials per repetition). None of the words used in this procedure were part of the stimulus set for the word categorisation task. The stimuli were generated with a text-to-speech software provided by Microsoft Cognitive Services (https://microsoft.com/cognitive-services/text-to-speech/#features, retrieved 2016/2017), using the male German (Austria) voice.

***Selection and validation of the stimulus material***

*Selection of stimulus material.* Using the Edinburgh associative thesaurus (Kiss et al., 1973) a pool of words associated with the target terms "sleep" and "activity" was generated. All words that appeared in both lists, consisted of more than one word, were antonyms to the target term, had the same word stem as the target term, or were not easily translatable to German, were removed. Each list was sorted according to the strength of association with the target term. The first 42 words of each list (sleep and activity), together with 42 words related to the target term "family", were presented to a total of N = 134 subjects (103 females and 31 males, 18 – 64 years old (median: 24)) in an online survey created using the platform SoSci Survey (Leiner, 2016). Subjects of the online survey rated all words of one list, intermixed with words of the two other lists. They rated valence, arousal, imaginability and how strong they associated the words with the target term (target terms being sleep (n = 50), activity (n = 36), or family (n = 48)). Using the results of this survey, we calculated sensitivity (affiliation of each word to its corresponding list) and specificity (demarcation of each word from the other two lists). We then chose 18 words of each list, parallelising them according to valence, imaginability, sensitivity, specificity, amount of syllables and word frequency in German (Quasthoff et al., 2011). After a piloting of the here presented experiment, the word pool was further reduced to 9 words for each list, removing words that showed to be problematic in the pilot and adding 4 words for reasons of face validity ("to lie down", "to stand", "deep sleep", and "high performance"). The final word list can be found in Table 1 of the main manuscript and the stimuli characteristics can be found in Table S2 in the supplement.

*Validation of stimulus material.* The stimuli used in the word categorisation task were validated once more in the main experiment. All 18 stimuli from the word categorisation task were visually presented in a randomized order while participants were sitting, after having performed the word categorisation task. For each stimulus participants rated which category (sleep vs. activity) the word belonged to based on their personal preference. Additionally, they rated valence and arousal on a five-point SAM scale as well as imaginability on a four-point scale from “not at all” to “very good”. There was no time restriction for responding and subjects could revise their responses. Sleep- and activity-related words did not differ in valence and imaginability. Activity-related words were much more arousing than sleep-related words. Sleep-related words were falsely categorised slightly more often compared to activity-related words. However, this effect was rather small and words were rarely miscategorised at all.

Table S2: Mean ± SD of values for valence, arousal, and imaginability for sleep- and activity-related words. Categorisation errors represent the words that were categorised into the opposite list as intended by the experimenters (e.g. categorising a sleep-related word as activity-related). Valence and arousal were rated on a five-point SAM scale (valence: 1 = “negative”, 5 = “positive”; arousal: 1 = “calm”, 5 = “arousing”). Imaginability was rated on a four-point scale (1 = “not at all”, 4 = “very good”).

| | **sleep words** | **activity words** | **Statistics** |
|---|---|---|---|
| **valence** | 3.45 ± .36 | 3.45 ± .42 | $T(65) = .13$, $p = .90$ |
| **arousal** | 1.90 ± .43 | 2.96 ± .73 | $T(65) = 13.36$, $p < .01$, $\omega^2 = .57$ |
| **imaginability** | 3.38 ± .42 | 3.32 ± .43 | $T(65) = -1.66$, $p = .10$ |
| **categorisation error (%)** | 3.37 ± 7.54 | 1.18 ± 4.83 | $T(65) = -2.85$, $p < .01$, $\omega^2 = .05$ |

***Volume effects on the event-related potentials***

N180 and P280 were strongly influenced by volume. The mean amplitude of both components increases with stimulus intensity, as it is typical for early ERP components reflecting the physical properties of stimuli. For the N180, the effect spanned both hemispheres, but was largest in the left hemisphere. The contributing electrodes formed a single significant cluster (F-mass: 361.21, $p < .01$; cluster size: 38, $p < .01$). Effect sizes for the contributing electrodes ranged between $.02 < \omega^2 < .20$, with an average (± SD) of $\omega^2 = .09 \pm .05$. The effect was maximal at CP3 ($F(3,195) = 22.51$, $p < .01$, $\omega^2 = .20$) and had the same direction for all significant electrodes (see Figure S5, outer left columns).

Post hoc t-tests for the N180 revealed that mean amplitude of the EEG components differed significantly and strongly (according to the effect size $\omega^2$) between all volume levels, except for the two intermediate volume levels (-.05 vs. ±0 dB SPL). The difference was evident most strongly in the parietal cortex. The post hoc t-tests had a minimum of 95 % power (calculated individually for each t-test and electrode) to find effect sizes ($\Omega^2$) of 0.05 ± 0.01 (range: 0.02 - 0.09) or larger for the N180.

Similarly, for the P280 the effect of volume on mean amplitude spanned both hemispheres. However, for the P280 the effect was largest in the right hemisphere. A significant cluster was formed by fronto-central electrodes (F-mass: 77.51, $p < .01$; cluster size: 14, $p < .01$) Effect sizes for the contributing electrodes ranged between $.02 < \omega^2 < .12$, with an average (± SD) of $\omega^2 = .05 \pm .03$. The effect was maximal at FCz ($F(3,195) = 13.09$, $p < .01$, $\omega^2 = .12$) and had the same direction for all significant electrodes (see Figure S5, inner left columns). The effect was reflected as negativity in the parieto-occipital cortex; however these electrodes did not form a significant cluster (F-mass: 26.44, $p = .05$; cluster size: 6, $p = .06$).

Post hoc t-tests for the P280 revealed that mean amplitude of the EEG components differed significantly and strongly (according to the effect size $\omega^2$) between the lowest volume level (-1.0 dB SPL) and the highest volume level (+0.5 dB SPL). While the effect sizes, i.e. the differences in mean amplitude between volume levels, became smaller when comparing the lowest against the second loudest as well as the second lowest against the loudest volume level. Effect sizes decreased even further when comparing directly neighbouring volume levels (-1.0 vs -0.5 dB SPL, -0.5 vs ±0 dB SPL, and ±0 vs +0.5 dB SPL). Power for the post hoc t-tests is lower than for the ANOVA, and differences (reflected in smaller effect sizes) in mean amplitude were smaller the closer the analysed volume levels were. Therefore, post hoc t-tests comparing directly neighbouring volume levels mostly showed non-significant results. The post hoc t-tests had a minimum of 95 % power (calculated individually for each t-test and electrode) to find effect sizes ($\Omega^2$) of 0.04 ± 0.01 (range: 0.02 - 0.06) or larger for the P280.

Likewise, volume influenced the N400 globally. Its effect showed up in two separate clusters: an early cluster (400-900 ms; F-mass: 224.36, $p < .01$; cluster size: 49, $p < .01$) and a late cluster (900-1700; F-mass: 986.25, $p < .01$; cluster size: 149, $p < .01$) (see Figure S5, right columns). Effect sizes for the contributing electrodes in the early cluster ranged between $.02 < \omega^2 < .09$, with an average (± SD) of $\omega^2 = .04 \pm .02$. The effect was maximal in C1 in the time window 400-500 ms for the early cluster ($F(3,195) = 9.50$, $p < .01$, $\omega^2 = .09$). In the early time window, the N400 was less pronounced for the lowest volume level compared to all other volume levels. Effect sizes for the contributing electrodes in the late cluster ranged between $.02 < \omega^2 < .17$, with an average (± SD) of $\omega^2 = .06 \pm .04$. The effect was maximal in P1 in the time window 1100-1200 ms for the late cluster ($F(3,195) = 18.69$, $p < .01$, $\omega^2 = .17$). The volume effect on the N400 was maximal in the time window from 1000 to 1400 ms and had the same direction for all significant electrodes and time windows. In the late cluster, in contrast to the early components (N180 and P280), lower volumes led to a higher mean activity compared to higher volumes. This indicates that it is more difficult and takes longer to infer the meaning of a word presented at volumes below the speech recognition threshold.

Post hoc t-tests for the N400 revealed that mean amplitude of the EEG components differed significantly and strongly (according to the effect size $\omega^2$) between the lowest volume level (-1.0 dB SPL) and all other volume levels. The three other volume levels themselves barely showed significant differences in mean amplitude when compared with each other. The post hoc t-tests had a minimum of 95 % power (calculated individually for each t-test and electrode) to find effect sizes ($\Omega^2$) of 0.03 ± 0.01 (range: 0.01 - 0.08) or larger for the N400.

Descriptively the lowest volume always had the lowest mean activity for the N180 and P280, with increasing mean amplitude as volume increased. In contrast, for the N400, mean amplitude was largest for the lowest volume in the late cluster and significantly lower in all other volume levels.

The ANOVA itself had a minimum of 95 % power to find effect sizes ($\Omega^2$, calculated individually for each electrode) of .03 ± .01 (range: .03 - .05) or larger for the N180, of .03 ± .00 (range: .02 - .03) or larger for the P280, and .02 ± .01 (range: .01 - .05) or larger for the N400.

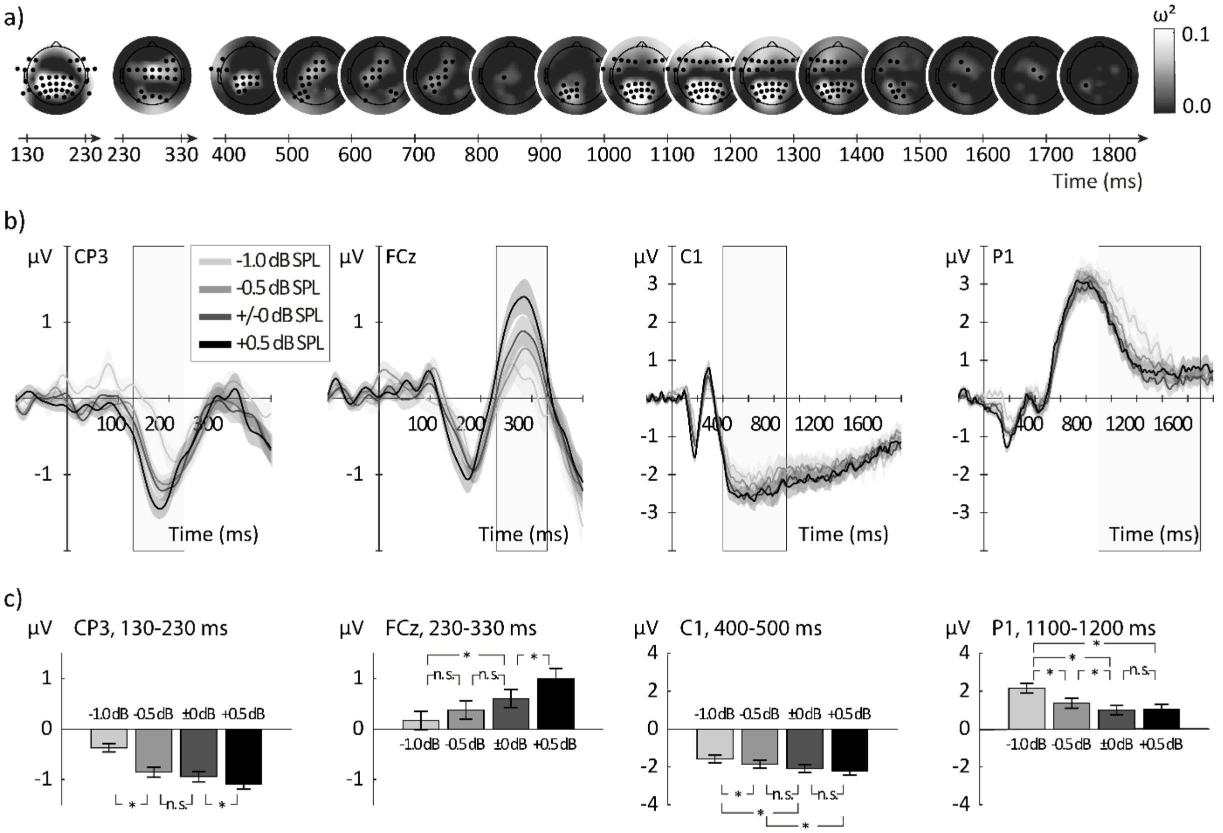


Figure S5: a) Topographic maps of the effect sizes ($\omega^2$) from the main effect volume of the ANOVA (comparison of the four volume levels averaged over congruent and incongruent trials). Electrodes showing significant differences are plotted as black dots. Clusters for the volume effect span large parts of the cortex. For the N400 two distinct clusters are found, an early left-lateralised cluster and a late cluster over the frontal and parietal cortex. Lighter colours indicate larger effect sizes. b) Grand average event-related potentials (± SE) averaged over all participants and congruency conditions, plotted separately for the four volume levels (from light grey [-1.0 dB SPL] to black [+0.5 dB SPL]). Exemplary electrodes from the N180 cluster (CP3), from the P280 cluster (FCz), from the early N400 cluster (C1), and from the late N400 cluster (P1) are depicted. The grey rectangle marks the period with significant differences between the volume levels. For the N180 and the P280 the mean amplitude increased with increasing volume levels. Similarly, the early N400 cluster revealed that the lowest volume level exhibited significantly less pronounced mean amplitude compared to the three other volume levels, which do not differ between each other. In contrast, for the late N400 cluster, mean amplitude was largest for the lowest volume and significantly lower in all other volume levels. c) Mean amplitude (± SE) for each volume level averaged over all participants and congruency conditions. The chosen electrodes and the chosen time windows reflect the periods where the effect was maximal for each cluster. All electrodes contributing to a cluster had the same effect pattern as depicted in these exemplary electrodes. While amplitude significantly decreases (N180) or increases (P280) with increasing volume levels for the early components, in the N400 only the lowest volume level differs significantly from all other, which themselves do not exhibit significant amplitude differences. *: $p < .05$.
